# Diverse strategies for tracking seasonal environmental niches at hemispheric scale

**DOI:** 10.1101/2022.07.15.500241

**Authors:** Jeremy Cohen, Walter Jetz

**Affiliations:** Department of Ecology and Evolutionary Biology, Yale University, 165 Prospect Street, New Haven, CT, 06520 USA; Center for Biodiversity and Global Change, Yale University, 165 Prospect Street, New Haven, CT, 06520 USA

**Keywords:** Environmental niche, climatic niche, hypervolume, seasonal niche tracking, birds, big data, seasonality, migration, functional trait groups, phylogenetic signal, GBIF

## Abstract

Species depend upon a constrained set of environmental conditions, or niches, for survival and reproduction that are increasingly lost under climatic change. Seasonal environments require species to either track their niches via movement or undergo physiological or behavioral changes to survive. Here we identify the tracking of both environmental niche position and breadth across 619 New World bird species and assess their phylogenetic and functional underpinning. Partitioning niche position and breadth tracking can inform whether climatic means or extremes limit seasonal distributions. We uncover diverse strategies, including the tracking of niche position, breadth, both, or neither, suggesting highly variable sensitivity to ongoing climatic change. There was limited phylogenetic determinism to this variation, but a strong association with functional attributes that differed between niche position and breadth tracking. Our findings imply significant functional consequences for communities and ecosystems as impending climate change affects some niche tracking strategies more than others.

## Introduction

Species survive and reproduce under a specific set of environmental conditions, known as the environmental niche or *n*-dimensional hypervolume (Blonder et al. 2014; Hutchinson 1957; Lu, Winner, and Jetz 2021). In seasonal environments, species must adjust to a constantly shifting window of available conditions through one of several strategies. When remaining stationary, species must either maintain tolerance to a wide range of conditions or undergo physiological and behavioral changes to survive seasonal variation, known as ‘niche switching’ (Nakazawa et al. 2004). Alternatively, seasonally mobile animals can occupy a dynamic niche that remains relatively narrow across the annual cycle, known as ‘niche tracking’ (Gómez et al. 2016; Somveille, Rodrigues, and Manica 2018; Winger et al. 2019). Seasonal niche tracking is central to the persistence of species with limited behavioral or physiological capacity to adjust their niches (Fandos et al. 2020; Zurell et al. 2018). Understanding the functional and phylogenetic drivers of niche tracking behavior across diverse species can allow researchers to predict how species mediate exposure to novel, potentially adverse conditions as climate change progresses (Tingley et al. 2009; La Sorte and Jetz 2012; Somveille, Rodrigues, and Manica 2015).

After the documentation of seasonal niche tracking behavior in single species (Fandos et al. 2020) and smaller clades (Gómez et al. 2016; Eyres et al. 2020), a more general understanding of the patterns, causes and consequences of seasonal niche tracking across diverse taxonomic groups remains missing. For example, the role of phylogeny and functional traits in driving seasonal niche tracking across a diverse species set remains largely unexplored (but see Zurell et al. 2018). Several studies have hypothesized that niche tracking may be phylogenetically conserved (Gómez et al. 2016; Martínez–Meyer, Townsend Peterson, and Navarro–Sigüenza 2004), as is typical of behavioral and migratory traits (Outlaw and Voelker 2006), but this has not been evaluated rigorously or broadly. Alternatively, niche tracking may have repeatedly evolved in tandem with species’ functional traits. For example, obligate insectivores or small-bodied species may be most likely to closely track their niche over the annual cycle to satisfy narrow dietary or thermal requirements (Gómez et al. 2016; Huey et al. 2012). Separating these potential drivers will allow researchers to better predict niche flexibility and niche tracking, and thus climate change vulnerability, among rare species or those from under-sampled regions.

Previous work on niche tracking has focused on seasonal similarity in niche positions, or mean environmental conditions, without accounting for niche components such as niche breadth (the volume or range of tolerable conditions). However, niche breadth is a central additional dimension because climate change is altering both the means and variances of climatic conditions (Rahmstorf and Coumou 2011). For example, species may seasonally track niche breadth instead of changing the central niche position when extremes are more limiting to their survival and reproductive success than climatic means, as is the case for numerous species (Albright et al. 2010; Ummenhofer and Meehl 2017). Thus, species may use the tracking of niche breadth as an alternative strategy to niche position tracking to persist in the face of seasonality, one unexplored by the existing niche tracking literature, which instead considers niche breadth as an annually static variable (Gómez et al. 2016; Zurell et al. 2018). To better identify complex variation in niche tracking, seasonal niche similarity should be partitioned into constituent components, including *niche position similarity*, or the distance between niche centroids, representing the difference between the average conditions a species experiences in each season; and *niche breadth similarity*, or the proportional difference in niche breadths (Lu, Winner, and Jetz 2021), representing the range of conditions a species can tolerate. Hypervolumes quantified using non-parametric techniques, such as kernel-density estimates or support vector machines (Blonder 2018; Brown, Holland, and Jordan 2020), are difficult to partition into constituent components. However, recently developed parametric methods for quantifying the niche allow for partitioning of these components and direct hypothesis testing against predictions derived from theories (Lu, Winner, and Jetz 2021).

We use these new metrics to distinguish five primary strategies for seasonal niche tracking (Fig. 1; Fig. 2): i) “Non-trackers” retain neither the position nor the breadth of their niches over the annual cycle; ii) “Breadth trackers”, or “position shifters”, track niche breadth but shift their position, suggesting that their seasonal ranges may be limited by environmental variation; iii) “Position trackers”, or “breadth expanders”, track only niche position and adjust niche breadth between seasons and may be seasonally limited by environmental means; Finally, “complete trackers” track both niche position and breadth, but may do so by either iv) migrating to track weather conditions across the annual cycle or v) remaining stationary in an aseasonal environment.

**Figure 1:**
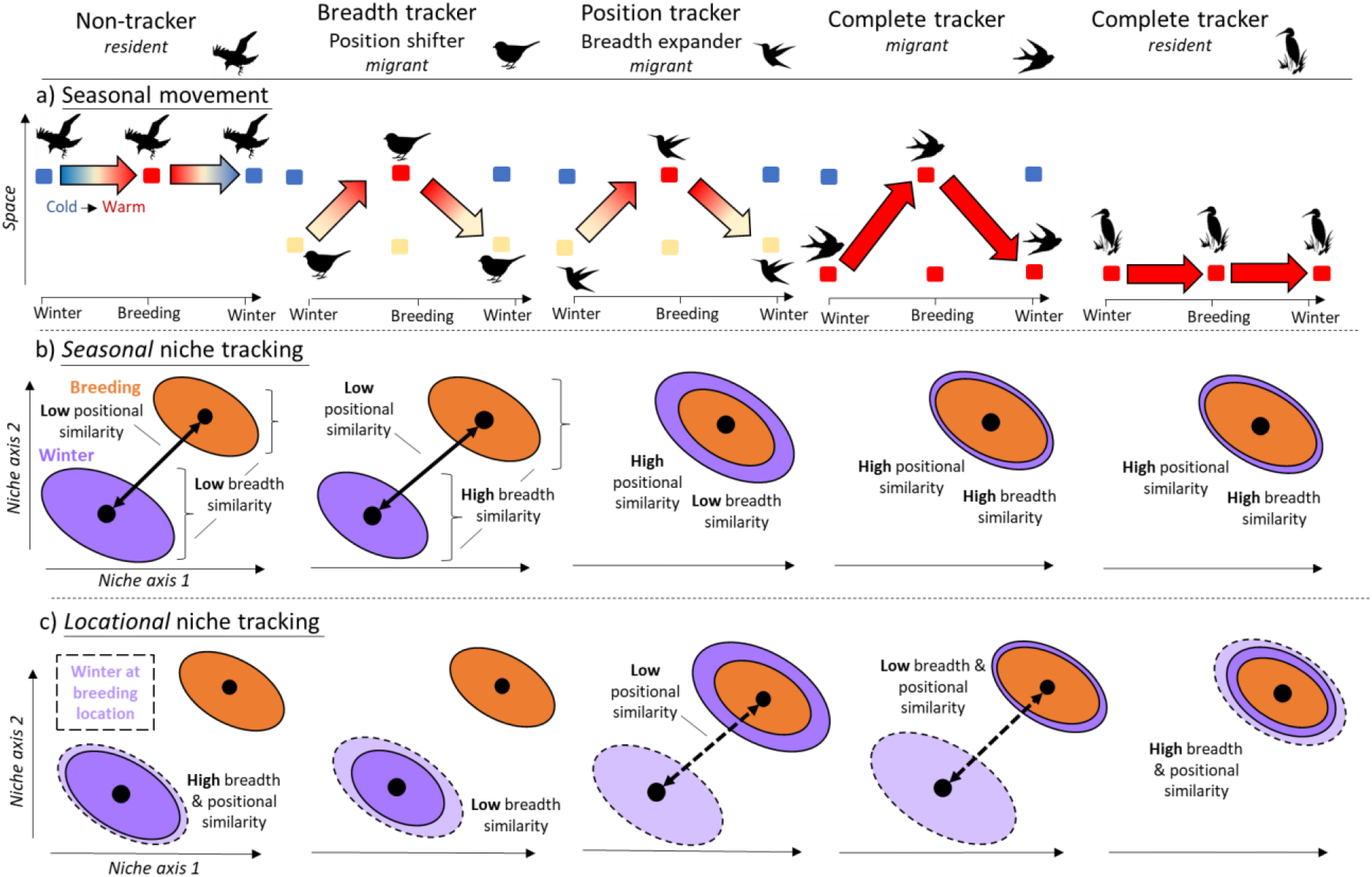
A typology of seasonal niche tracking strategies. Seasonal niche similarity is decomposed into two components, similarity in niche position and niche breadth. A given species may (1) be a non-tracker, tracking neither niche position or breadth, (2) track niche breadth, but shift position, (3) track niche position, but expand breadth, or (4-5), be a complete tracker, tracking both niche position and breadth. a) shows the differing use of climate zones over space and time; b) compares seasonal niches between the breeding (orange) and overwintering (purple) seasons (solid arrows represent positional similarity, and brackets represent breadth similarity); c) compares the location of the winter conditions at breeding locations (light purple) and the overwintering niche, revealing that for strategy 4 and 5 niche tracking is achievable through multiple life history strategies. Dashed arrows represent locational positional similarity.

**Figure 2.**
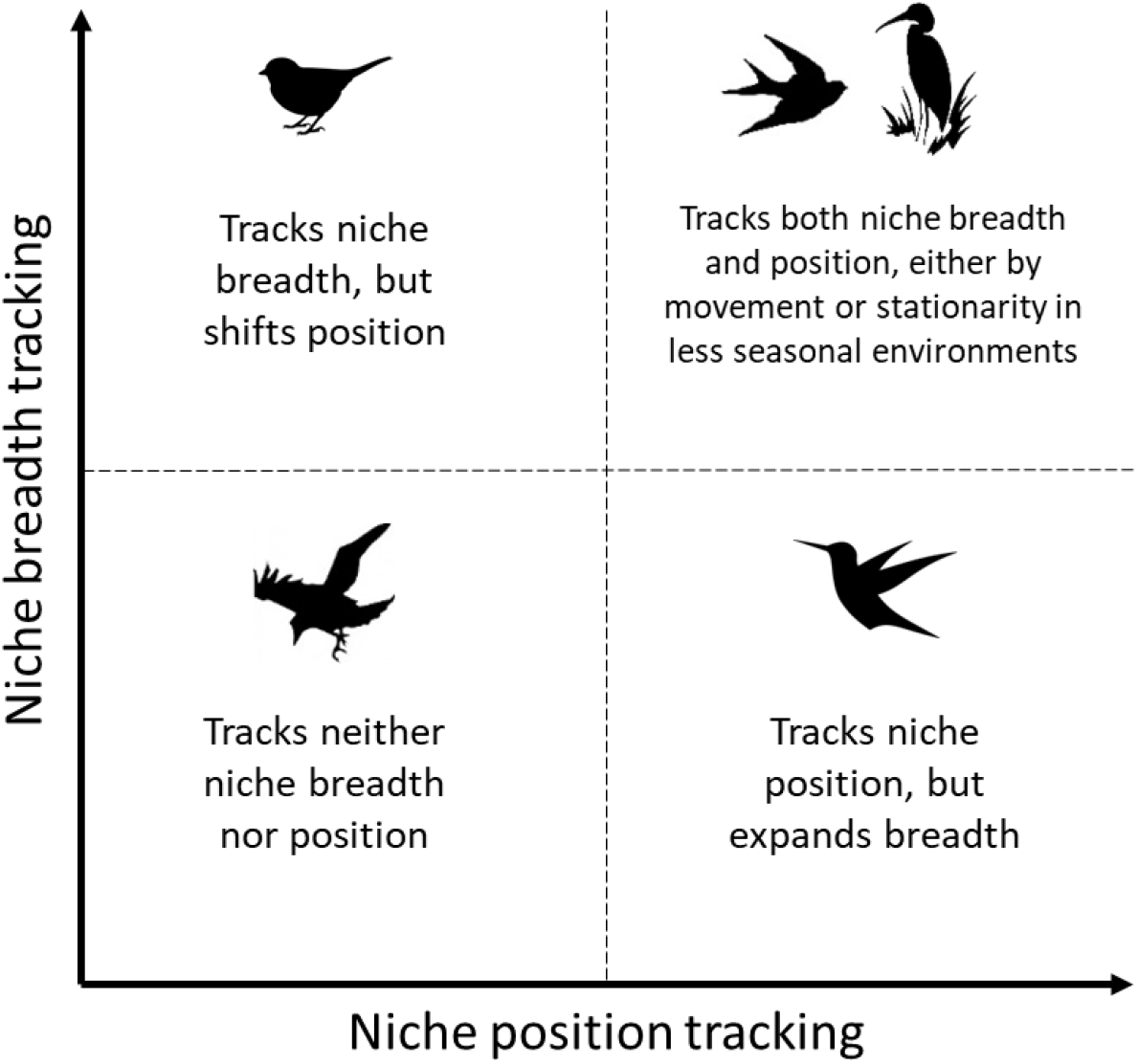
Five niche tracking strategies. Conceptual schematic outlining the contribution of niche breadth and position tracking to each of five possible niche tracking strategies.

However, a simple comparison of niche space during the breeding and overwintering seasons (henceforth, *seasonal* niche similarity) cannot fully distinguish these strategies (Fig.1b, columns 4-5), despite obviously divergent dispersal capability and potential for climate change adaptation (Eyres et al. 2020). We therefore consider an additional metric, *locational* niche similarity, in which the realized overwintering hypervolume is compared with that had the species remained at its breeding range, its *stationary winter niche* (Fig. 1c). This measure, which emphasizes the similarity of conditions at specific locations, is important for understanding seasonal niche tracking because it accounts for the environmental distance covered by the species through purposeful movement. Among temperate breeders, seasonal and locational niche similarity are likely to have an inverse relationship; for example, a niche tracking species with little difference between its breeding and overwintering niche is likely to experience highly distinct overwintering conditions compared with those at its breeding sites.

Here, we use a hemispheric system to address patterns and drivers of these strategies; specifically, 619 bird species, representing nearly the full diversity of birds breeding in the US and Canada. This species set is ideally suited given its tremendous variation in migratory strategy, diversity of functional trait groups (Barnagaud et al. 2017) and uniquely comprehensive occurrence data. We also leverage a recently developed environmental annotation tool (Li et al. 2021) and a novel parametric hypervolume method (Lu, Winner, and Jetz 2021) to quantify similarity between the environmental niches occupied by each species during both the breeding and overwintering seasons. We use this system to address the following questions:

1. What is the prevalence of different niche tracking strategies while accounting for both niche position and breadth? To date, a quantitative multi-species assessment of strategies across a diverse taxonomic group has been missing.
2. What is the relationship between the tracking of niche position and the tracking of niche breadth? Do species that maximize niche positional similarity across seasons also tend to retain similar niche breadth, or do species largely track only one or the other?
3. How phylogenetically and functionally determined are niche tracking strategies? Across functional trait groups, we hypothesize seasonal niche tracking to be most common in a) long-distance migrants, because they can physically relocate to suitable locations (Laube, Graham, and Böhning-Gaese 2015; Zurell et al. 2018); b) small-bodied species, because they have low thermal inertia and generally narrow thermal breadths (Huey et al. 2012; Albright et al. 2017), suggesting they cannot tolerate large seasonal variation in the niche; c) insectivores, because abundant insect prey is only available under specific temperature, precipitation and productivity levels (Winkler, Luo, and Rakhimberdiev 2013); and d) species occupying open or water habitats, because they are not shielded from climate variability by forest structure (Jarzyna et al. 2016). Given that behavioral and migratory traits are often phylogenetically conserved (Outlaw and Voelker 2006), we expect a strong phylogenetic signal in seasonal niche tracking behavior (Gómez et al. 2016).
4. Do cross-species comparisons of *locational* niche similarity reveal important behavioral strategies and functional or phylogenetic associations not apparent when quantifying only *seasonal* niche similarity? We predict an inverse relationship between seasonal and locational similarity across species and expect to observe functional trait relationships with locational similarity that are not observed with temporal similarity.

We expect the emerging insights to not only address these questions but more generally offer an assessment of seasonal niche dynamics as a system to understand realized strategies for mitigating exposure to climatic change.

## Material and Methods

### Species selection and environmental data

We identified the 672 bird species that annually breed or overwinter in the United States and Canada based on American Birding Association birding codes (Association 2008) updated to the Clements bird taxonomy as of 2021 (Clements 2007). These codes are a widely-accepted authority to distinguish regularly occurring species from irregularly occurring vagrants, or those species which sporadically appear on the continent each year but whose occurrence is not predictably tied to a given location. We excluded species not native to the US or Canada or those that are primarily marine, for which weather data is unavailable.

In August 2021, we accessed the Spatiotemporal Observation Annotation Tool (STOAT) v1.0, a novel cloud-based toolbox for flexible biodiversity annotations (Li et al. 2021), to download annotated Global Biodiversity Information Facility data (https://www.gbif.org/) for all species (data compilation, analyses and visualizations were all completed in R 4.1.0; R Core Team 2021). To minimize the potential for spatiotemporal sampling bias, we then thinned points by selecting one point from each location (5×5km grid cell) per week.

We annotated observation points with three environmental dimensions: daily maximum temperature (sourced from NASA-MODIS; https://lpdaac.usgs.gov/products/mod11a1v006/), enhanced vegetation index (EVI; from MODIS), and precipitation (from CHELSA v2.1; Karger et al. 2021), each summarized to a 1km buffer (0.5km radius) over 30 days prior to the observation (imported using jsonlite and httr packages; Ooms 2014; Wickham and Wickham 2020). We did not assess *a priori* whether these variables are equally relevant across species; for example, certain species may be limited by precipitation but not temperature within their range, or *vice versa*. Alternatively, some species may be limited by environmental factors not considered in our estimation of species niches. However, this variable set represents the environmental factors that most commonly drive species distributions (Qian 2010), and these variables have low collinearity, allowing each axis to remain independent. Estimating niches with the same variable set for all species was necessary to ensure consistency in cross-species evaluation of seasonal changes to both niche position and breadth.

For each species, we restricted observation points to those with environmental data available for all dimensions. Further, we temporally cropped data to season (December-February for the overwintering season and June-August for breeding season) and spatially cropped points to the American continents (< -30° longitude). Sufficient data for analysis (>20 points per season after filtering) was available for 619 species. Database management was completed using *tidyverse* packages (Wickham et al. 2019).

To quantify locational similarity, we conducted the same niche characterizations for breeding locations during the overwintering season. For each species, we created a hypothetical set of occurrence points corresponding to coordinate locations during the breeding season, each with randomized winter calendar dates (December-February), annotating them with environmental data as described above.

### Ecological niche modeling and seasonal similarity

For every species, we calculated parametric measurements of the similarity between the seasonal three-dimensional hypervolumes. The hypervolumes were characterized as multivariate normal distributions, allowing us to derive analytical estimates for the breadth and position of each hypervolume (MVNH package; Lu, Winner, and Jetz 2021). We partitioned niche similarity into two metrics: i) *niche position similarity*, which quantifies the distance among hypervolume centroids in each season based on the sign-flipped, log-transformed Mahalanobis distance, used to assess changes in niche position between seasons; and ii) *niche breadth similarity*, which represents the similarity in *niche breadth* (volume) between seasons and is measured as the sign-flipped, log-transformed determinant ratio.

To quantify seasonal similarity, we calculated positional and breadth similarity between the breeding and overwintering seasonal hypervolumes. To quantify locational similarity, we calculated positional similarity between the overwintering hypervolume and available hypervolume during winter at breeding locations. Thus, we had a total of three metrics of seasonal niche similarity per species. We visualized pairwise two-dimensional hypervolumes for each species using *ggplot2* (Wickham 2011).

### Functional traits and phylogeny

We obtained species-level functional trait values and an avian phylogenetic tree to test associations between trait groups, phylogeny, and seasonal niche tracking. We derived migration distances from (La Sorte et al. 2022; La Sorte personal communication), body mass from the Eltonian trait database (Wilman et al. 2014), and habitat preference and diet from Barnagaud et al. (2017). We grouped several categories of each categorical predictor to avoid false positives associated with small sample sizes and to keep our conclusions broad. Diet categories included carnivore, invertebrate, omnivore, and herbivore (combining ‘fruit’, ‘nectar’, ‘vegetation’, and ‘seed’ categories). Habitat categories included water (‘coastal’, ‘open_water’, ‘riparian_wetlands’), open (‘semi-open’, ‘rock’, ‘arid’), generalist (‘urban’, ‘developed’), and forest.

We updated an avian phylogeny from Jetz et al. (2012) to account for recent taxonomic changes, updating species names to the Clements bird taxonomy as of 2021 (Clements 2007) and treating recently split species as having no phylogenetic distance (Appendix 2). We also harmonized species names in all trait datasets to Clements.

### Cross-species models

To assess the role of functional traits in seasonal niche tracking, we used weighted multivariate phylogenetic generalized least squares (PGLS) models (caper package; Orme et al. 2013).

Phylogenetically correlated model errors in PGLS account for the non-independence of the species due to their phylogenetic relatedness (Symonds and Blomberg 2014). The dependence of the model errors arises from trait axes that we did not include in the analysis and that may be subject to niche conservatism so that model errors reflect the unobserved trait and thus the phylogenetic distance between species. We used species’ similarity in niche position and breadth as the response variables in the models and explained their variation with categorical trait variables. Prior to use in models, continuous predictor variables (migration distance and log-transformed body mass) were scaled to improve model fit and response variables were log-transformed and sign-flipped (representing niche similarity rather than dissimilarity) to improve interpretability. We weighted all species points in models by log-transformed minimum sample size (number of points in the season with less data) to account for uncertainty in seasonal niche dissimilarity estimates (true error estimates are unavailable when estimating niche dissimilarity). We fit three models testing the relationships between the four functional traits and three metrics of niche similarity: seasonal niche position similarity, seasonal niche breadth similarity, and locational niche position similarity. To evaluate the reliability of multivariate models, we also fit univariate models for each combination of functional trait predictor and response variable.

To measure the role of phylogeny in both positional and breadth similarity, we quantified Blomberg’s K (Blomberg, Garland Jr et al. 2003) and compared it to a null distribution of K after randomizing species’ responses 1,000 times (picante package; Kembel et al. 2010). K < 1 suggests greater than expected phylogenetically-correlated variance within clades, while K > 1 suggests variance among clades (i.e., phylogenetically-conserved trait). We also quantified lambda to assess the extent of phylogenetic conservation of niche metrics via Brownian motion. We visualized relationships between niche metrics and functional traits using *ggplot2* (Wickham 2011), *ggExtra* (Attali and Baker 2019) and *RcolorBrewer* (Neuwirth and Neuwirth 2011). We visualized partial residuals using *visreg* (Breheny, Burchett, and Breheny 2020).

## Results

We found all five niche-tracking strategies represented among North American birds (Fig 3). For example, Red-bellied woodpeckers (*Melanerpes carolinus*; Fig. 3a) are largely stationary birds occupying highly seasonal environments, with low seasonal similarity between their breeding and overwintering niches, but high locational similarity between their overwintering and stationary winter niches. In contrast, Pine grosbeaks (*Pinicola enucleator*; Fig. 3b) are short-distance migrants and “breadth trackers”, tracking niche breadth while shifting position across seasons. Migratory Swainson’s thrushes (*Catharus ustulatus*; Fig. 3c) are “position trackers”, tracking their niche position across seasons, but expanding their niche during the breeding season. Prothonotary warblers (*Protonotaria citrea*; Fig. 3d) are long-distance migrants that abandon breeding territories to closely track both niche components across seasons, resulting in high seasonal and low locational similarity. Finally, Great kiskadees (*Pitangus sulphuratus*; Fig. 3e) are stationary tropical residents with conserved niches across the annual cycle, resulting in both high seasonal and high locational similarity. We select these species for visualizations because they represent each strategy well in both two-dimensional niche space (temperature and EVI only; Fig. 3) and three-dimensional space (remainder of results).

**Figure 3:**
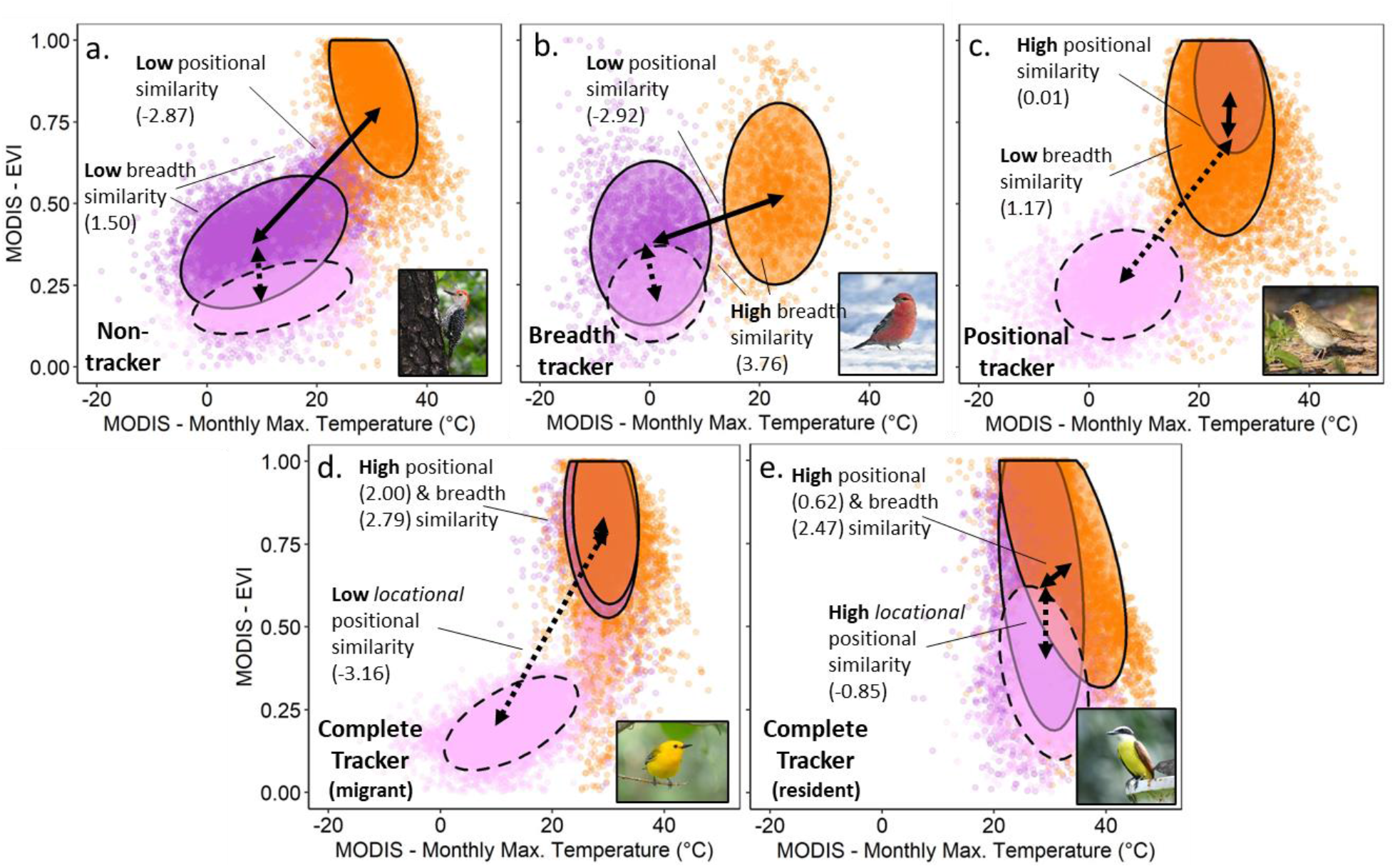
Diverse niche tracking strategies. (**a**) Red-bellied woodpeckers (*Melanerpes carolinus*) are non-trackers, with low *seasonal* similarity (solid arrow) between their breeding (orange points and ellipsoid) and overwintering (dark purple) two-dimensional niche (presented as standardized ellipse area). Meanwhile, the difference between their overwintering niche and winter conditions at breeding locations (light purple, dotted ellipsoid), or *locational* similarity (dotted arrow), is high. (**b**) Pine grosbeaks (*Pinicola enucleator*) track their niche breadth but shift position across seasons, while (**c**) Swainson’s thrushes (*Catharus ustulatus*) track their niche position but expand niche breadth. (**d**) Prothonotary warblers (*Protonotaria citrea*) closely track their niche across seasons via migration, resulting in high seasonal and low locational similarity. (**e**) Great kiskadees (*Pitangus sulphuratus*) track their niche by remaining stationary in less seasonal environments, leading to high seasonal and locational similarity. Position and breadth similarity values are provided in parentheses. Note that this figure displays two-dimensional niche space, though our statistical analyses consider three-dimensional niches, including precipitation as additional dimension.

The 619 bird species analyzed spanned the full diversity of niche tracking strategies (Fig. 4; Appendix 1). Across seasons, the median positional similarity (-log[Mahalonobis distance]) was -1.95 +/- 0.05 SE (i.e., centroids approximately 3 SDs apart), suggesting that many species are not closely tracking this niche attribute over the annual cycle (Ponti et al. 2020). For 60 species (9.7%), centroids were within 1 SD, suggesting highly similar niches in each season for this subset. The median among-season similarity in niche breadth was 2.17 +/- 0.03, equivalent to a 4:1 ratio in niche breadth between seasons, suggesting that many species expand their niche breadths as they traverse seasons. 129 species (22%) had a niche breadth similarity equivalent to a < 2:1 ratio, suggesting similarly-sized seasonal niches.

**Figure 4:**
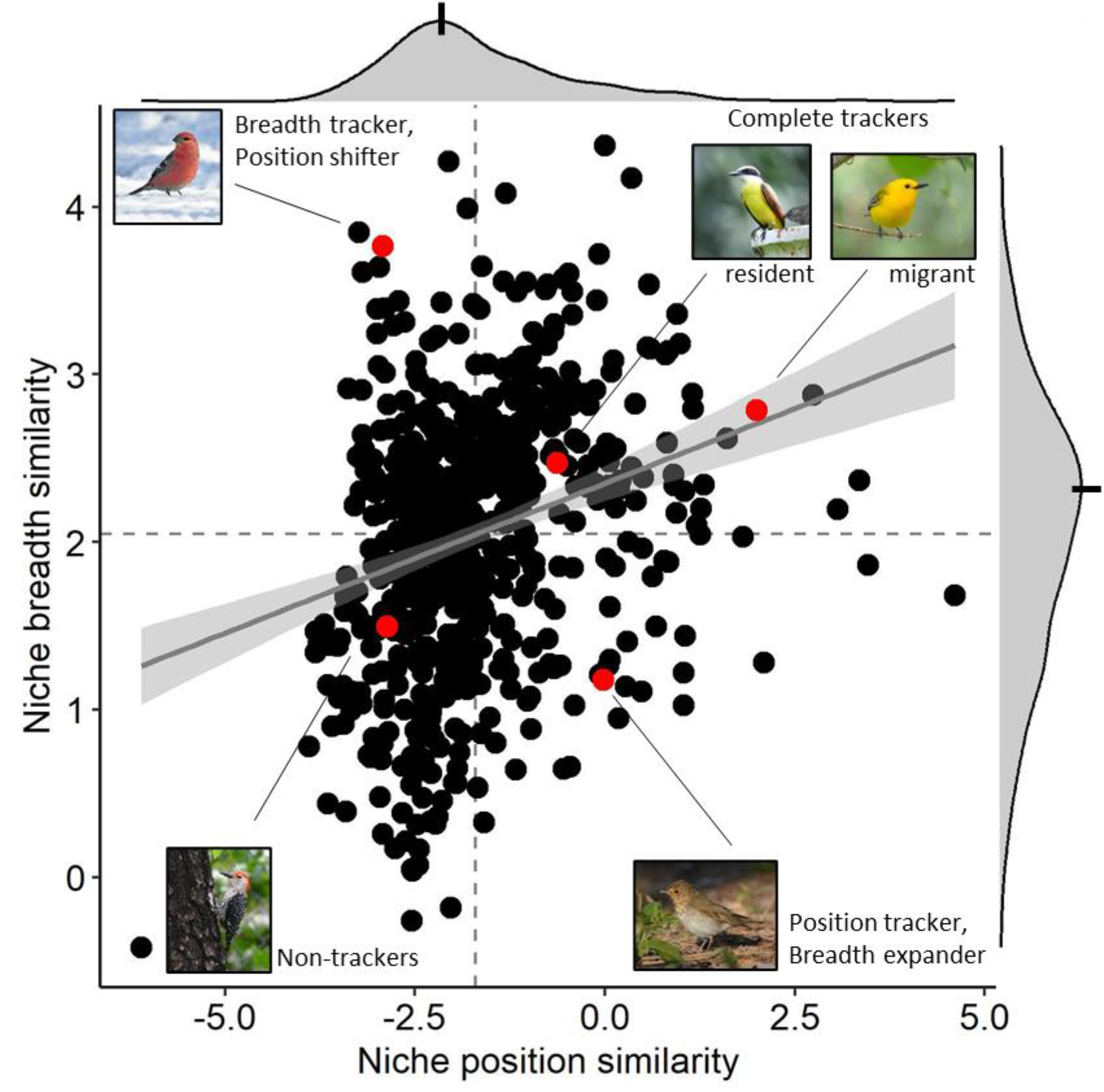
Interspecific variation in tracking niche position and breadth across seasons. The figure shows niche similarities between the breeding and overwintering seasons for the 619 bird species analyzed based on three niche dimensions (for red highlights, see Fig. 3). Strategies of tracking niche position, breadth, both, or neither are partitioned into four categories (boxes separated by dashed lines). Dashed lines highlight the means. Marginal density plots illustrate that species tracking only one niche component often track niche breadth more closely than niche position, with medians represented by notches. The trendline represents a linear relationship between niche position and breadth similarity with associated 95% confidence interval (shading). See Fig. S2 for equivalent patterns for locational similarity.

Tracking niche position was loosely tied to tracking niche breadth; species that tracked niche position more often tracked niche breadth, and *vice versa* (Generalized linear model: β=0.179, R^2^=0.073, p<0.001; Fig. 4). 170 (27.5%) species were “complete” niche trackers, tracking both position and breadth more than most species (positive values on both axes in Fig. 4), while 209 (33.8%) were non-trackers, tracking both less than others (negative values in Fig. 4). However, numerous species tracked one but not the other (Fig. 4), suggesting a complex diversity of approaches to seasonal niche tracking across bird species. Only 80 species (12.9%) tracked niche position but expanded breadth, while double that number – 160 (25.8%) – tracked breadth but shifted position.

We found seasonal niche tracking strategies to be closely associated with functional traits, and their role to be consistent across weighted multivariate models accounting for all traits and phylogenetic structure simultaneously (Fig. 5; Tables S1-S2) as well as univariate models (Tables S3-S4). As expected, both niche positional and breadth similarity were very strongly linked to migration distance (Phylogenetic generalized least-squares models: position, β=0.43, p<0.001; breadth, β=0.25, p<0.001), which was the strongest functional predictor; long-distance migrants were most likely to maximize similarity, while residents minimized similarity and others fell in the middle (Fig. 5a,d). Body mass emerged as another important functional predictor of seasonal niche tracking, especially with regards to positional similarity. Body mass negatively predicted niche position similarity (β=-0.80, p<0.001) and was positively associated with breadth similarity (β=0.53, p<0.001). Small-bodied birds maximized seasonal similarity, while large-bodied birds minimized it (Fig. 5b,e). Finally, both diet (F_1,3_=5.70, p<0.005) and habitat preference (F_1,3_=9.68, p<0.001) categories described variation in positional similarity, though only habitat preference drove breadth similarity (F_1,3_=10.15, p<0.005). As predicted, insectivores maximized positional and breadth similarity, herbivores minimized these, and omnivores fell in the middle (Fig. 5c,g). Waterbirds tracked their niches more closely than other species (Fig. 5d,h).

**Figure 5:**
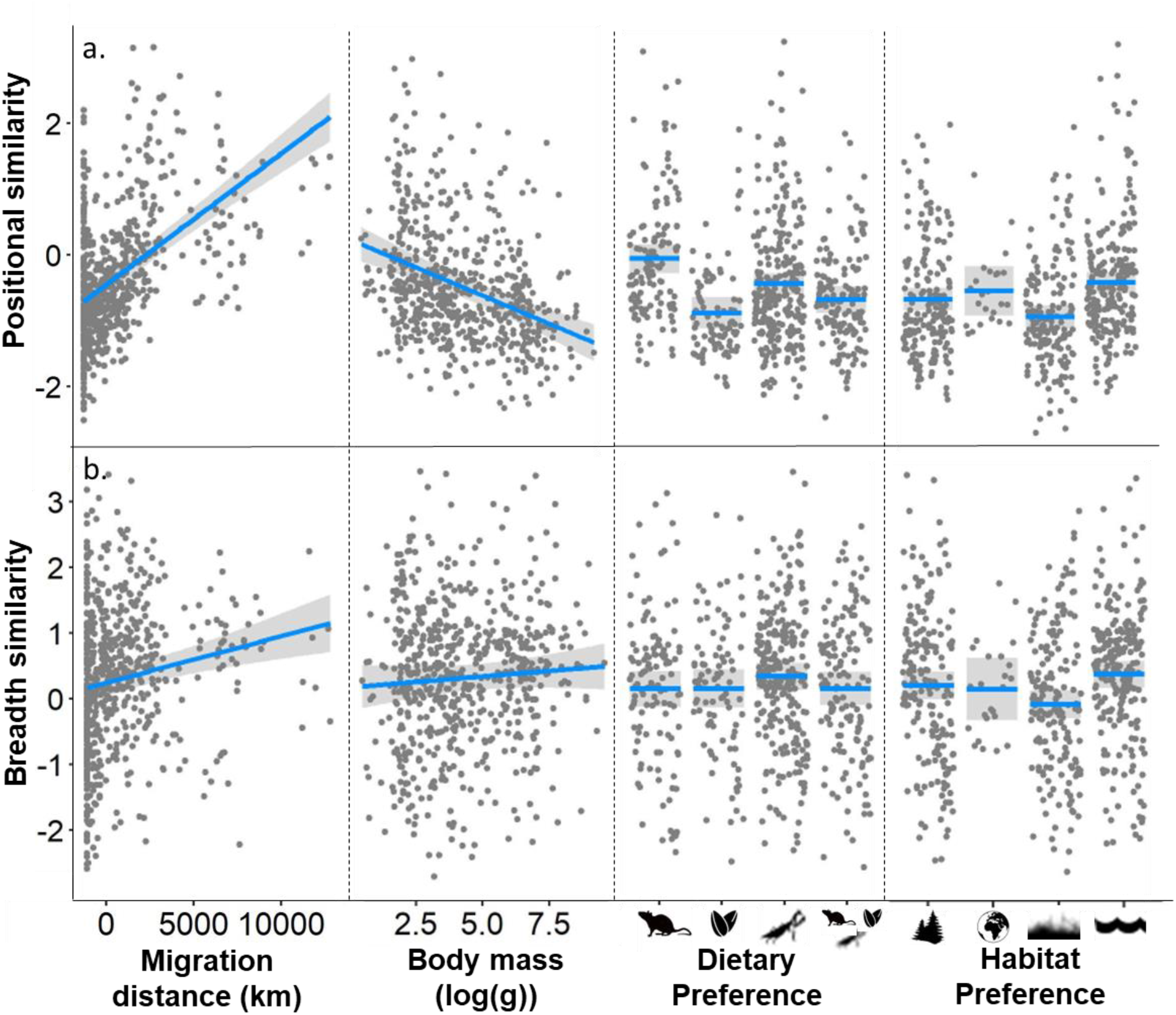
Functional traits predict variation in seasonal niche tracking, as shown by partial residual plots based on phylogenetic least-squares models. Y-axis values represent (a) positional and (b) breadth similarity. Silhouettes correspond to categorical functional trait values as follows: Dietary preference (left to right: carnivore, herbivore, insectivore, omnivore); Habitat preference (forest, generalist, open/grassland, water).

Surprisingly, we found little evidence of a phylogenetic signal in seasonal niche tracking. For both niche position and breadth, Within-group phylogenetic variation was greater and among-group variation lesser than expected by chance (Bloomberg’s K always < 0.1; position: p<0.001; breadth: p=0.10; Table S5) and did not support Ornstein-Uhlenbeck trait evolution. Measurements of lambda indicated a moderate phylogenetic signal for species tracking niche position (λ=0.66, p<0.001), suggesting possible Brownian-motion trait evolution, but little signal for niche breadth (λ=0.28, p<0.001).

Species that maximized seasonal niche similarity typically minimized locational niche similarity, and vice versa (β=-0.408, R^2^=0.237, p<0.001; Fig. S1). For locational similarity, the link between tracking niche position and niche breadth was stronger than for seasonal similarity, with 42% of species tracking both components and 29% neither (Fig. S2). Only 15% of species tracked locational niche position but not breadth, and 13% tracked locational breadth but not position. Likewise, relationships between functional traits and locational positional similarity were generally opposite to those observed with seasonal positional similarity. Long-distance migrants (β=-0.331, p<0.001), small-bodied birds (β=0.95, p<0.001), and waterbirds (F_1,3_=7.75, p<0.001) each minimized locational positional similarity, though we found no effect of diet. As with seasonal similarity, we found no evidence for a phylogenetic signal in locational niche similarity (Table S5).

## Discussion

For mobile organisms, seasonal niche tracking is a central part of the behavioral repertoire for mediating exposure to novel climate conditions. Here, we assessed the representation of distinct seasonal niche tracking strategies across North American breeding birds and evaluated the functional and phylogenetic drivers of these strategies. Our analyses revealed five distinct seasonal niche tracking strategies and their key functional drivers. We find that only a bit more than half of species can be categorized as either “complete trackers”, tracking both niche position and breadth across seasons, or “non-trackers”, tracking neither. However, almost 40% of species only track niche position or breadth, revealing complexity in niche tracking strategies that is previously unexplored in niche-tracking studies. Twice as many species tracked only niche breadth (26%) compared with those that tracked niche position and expanded niche breadth (13%), suggesting that tracking breadth is often a prerequisite for tracking position and that birds may be most specialized in terms of the range of environmental conditions they can tolerate rather than means. Importantly, niche tracking studies that do not consider niche breadth would likely have labeled the 26% of species that are niche breadth trackers as non-trackers, ignoring the role that niche breadth tracking and expansion plays in allowing species to persist over seasons. Species limited by extreme or highly variable weather conditions rather than climatic means may be more apt to prioritize niche breadth tracking while allowing niche position to seasonally fluctuate. Meanwhile, a species with inflexible thermal or dietary requirements that is specialized only during certain parts of the year (e.g., due to strict breeding requirements) may track niche position but not breadth. However, further work is needed to pinpoint the ecological motivations behind selection of breadth tracking as a niche tracking strategy.

We found that migration distance was the most important functional driver of seasonal niche tracking, despite this link previously receiving mixed support among cross-species niche tracking studies (Zurell et al. 2018; Laube, Graham, and Böhning-Gaese 2015; Gómez et al. 2016). Although several studies (Gómez et al. 2016; Laube, Graham, and Böhning-Gaese 2015) failed to detect a link between migration and niche tracking, their approaches compared species within single families, including residential species that are exclusively tropical. Because these residents occupy relatively stable environments, they can track their niches despite being stationary (Eyres et al. 2020). This example helps illustrate why migration distance and niche tracking are less correlated than might be expected – tropical residents can track their niches well despite their stationarity, while long-distance migrants may experience high seasonal variation despite their migration if they annually move between arctic breeding grounds and boreal or temperate wintering grounds. Indeed, although migration distance was the strongest correlate of seasonal niche tracking in our study, the correlation was moderate (R^2^=0.249).

Body size was revealed as the next most important functional trait associated with seasonal niche tracking when controlling for other traits. Small-bodied birds may not be able to tolerate large variation in their thermal niches because they have low thermal inertia (Huey et al. 2012; Albright et al. 2017). Further, small-bodied animals have shown greater sensitivity to extreme weather events (Cohen, Fink, and Zuckerberg 2021) and more rapidly alter phenological timing in tune with interannual weather variability (Cohen, Lajeunesse, and Rohr 2018) compared with large-bodied animals, suggesting that they must carefully maintain thermal limits. Additionally, small-bodied birds may be more likely to evolve niche-tracking strategies dependent on seasonal migration because long-distance flight carries lower energetic costs than it does for large birds (Watanabe 2016).

Finally, diet and habitat preference were additional meaningful drivers of seasonal niche tracking behavior, with insectivores and carnivores tracking their niches more than herbivores and omnivores, and waterbirds more so than other species. Insectivores may prioritize niche-tracking because they are dependent upon prey that is most commonly available under specific thermal and productivity limits (Winkler, Luo, and Rakhimberdiev 2013). Surprisingly, carnivores tracked their niche positions even more closely than insectivores, but many of these species are predating on herpetofauna also active during warm weather. Though we predicted both open and water habitat specialists to be the strongest niche trackers, as they are less shielded from the external environment by habitat structure (Jarzyna et al. 2016), only water birds tracked their niche more closely than other groups (Fig 4d,h). Bird species that occupy open habitats may make greater use of microhabitat structure than water birds, which are typically exposed (e.g., (Shew, Nielsen, and Sparling 2019)).

Although many studies have assumed seasonal niche tracking to be phylogenetically conserved (Gómez et al. 2016; Martínez–Meyer, Townsend Peterson, and Navarro–Sigüenza 2004), this has not to our knowledge been evaluated over a large, diverse taxonomic group such as North American breeding birds. There are several explanations as to why we did not detect a phylogenetic signal. First, migratory behavior has arisen in numerous, diverse lineages of birds, and is flexible even within species (Zink 2011). Second, closely related species often occupy distinct climatic zones that vary greatly in seasonality, especially if speciation resulted from character displacement (Newton 2003). Finally, species-level exposure to and adaptation potential for climate change may be quite independent from phylogeny (Khaliq et al. 2015; Davis et al. 2010). Based on our analyses, we conclude that functional traits likely play a larger role than phylogeny in driving seasonal niche tracking in birds.

Complex variation among niche tracking strategies cannot fully be assessed via seasonal niche comparisons; for example, a species that niche-tracks via movement and a stationary species in an aseasonal environment have similar seasonal niches. To parse these strategies, we estimated locational niche similarity, or the similarity between overwintering niches and the *stationary winter niche* for each species. Species with high seasonal similarity often had low locational similarity, and *vice versa*, reflecting a trade-off between conserving niches and ranges throughout the annual cycle. While many species tracked only niche position or breadth between seasons, species adhered to locationally tracking both or neither components much more often, with few tracking just one. Therefore, species may be leaving breeding areas in winter for the purposes of modifying both the mean and variance of environmental conditions. We observed several key differences between functional trait relationships and either seasonal or locational niche similarity. For instance, migration distance and body size were equally important as drivers of locational niche similarity, possibly because large-bodied birds are especially reluctant to energetically invest in movement (Watanabe 2016). Further, in contrast with seasonal niche similarity, there was no effect of diet on locational similarity, perhaps because food availability depends more on environmental conditions themselves rather than the distance birds travel to track them. Thus, we demonstrate the importance of considering both seasonal and locational similarity in species niches to understand variation in niche tracking strategies.

We estimated niche metrics at the species level to effectively compare niche tracking strategies across a taxonomic group as broad and diverse as birds and to make use of the enormous quantity of available species-level occurrence data. However, we did not explore niche variation as experienced at lower levels of organization, including populations, demographic groups, and individuals (Fandos et al. 2020; Fandos and Tellería 2020; Carlson et al. 2021). Populations of species with wide geographic ranges might be adapted to highly distinct environmental conditions (Broggi et al. 2005). Within a population, demographic groups can also occupy different niches – sexes can occupy spatially distinct regions during the nonbreeding season, or young can be shielded in microclimate during breeding season (Shipley et al. 2020). Finally, individuals may track different niche components than populations (Fandos et al. 2020) or might have strong variation in their ecological niches during different parts of the year and thus vary in niche-tracking. For example, individual white storks (*Ciconia ciconia*) and sandhill cranes (*Antigone canadensis*) range from subtropical residents to short-distance migrants to longer-distance migrants moving between northern temperate and tropical zones each year (Krapu et al. 2014; Fandos et al. 2020; Carlson et al. 2021). Summarizing niche tracking strategies at the species level may limit our inferential ability in two ways: 1) species-level niche breadth is likely greater than individual niche breadth given variability in individual niches (Carlson et al. 2021), and 2) the accuracy of species-level niche centroids for individuals likely depends on the variation in climate zones at which the species exists, so long as individuals and populations are locally adapted (Araújo and Costa-Pereira 2013). Although most occurrence data is still available at the species level, new technologies such as GPS tracking offer exciting opportunities for assessing individual niches over time and across hierarchies of organismal organization (Jetz et al. 2022; Costa-Pereira et al. 2022) and complement the presented species-level findings.

As climate change progresses, species are increasingly being exposed to conditions that are quite different from those to which they have adapted (Pacifici et al. 2015). Gradual shifts in range boundaries or phenology allow species to keep pace with long-term changes in climatic means (La Sorte and Jetz 2012; Rushing et al. 2020; Koleček, Adamík, and Reif 2020). However, niche tracking is an important tool for species to buffer both the changing mean environmental conditions and increasing interannual variability associated with climate change by selecting optimal environmental conditions over short time scales (Román-Palacios and Wiens 2020), especially for species with narrow thermal or habitat requirements. Our findings reveal a broad diversity in niche tracking strategies and uncover important functional trait associations with migratory behavior and body size. This has ecological consequences for the assemblages and ecosystems that will see different functional perturbations due to the different susceptibility of niche tracking strategies to climatic change (Barbet-Massin and Jetz 2015). For example, as niche tracking becomes more difficult with climate change, small-bodied and insectivorous birds may be especially likely to fail to maintain their niches, resulting in proportional declines of these species relative to the broader avian community assemblage.

Thus, we provide a framework for future studies to assess the degree of tracking both niche position and breadth and capability for climate change adaptation in many other species globally, including those from understudied regions with limited available data. However, additional work is needed to assess how species have already been keeping up with climate change that has occurred. Understanding how climate change is modifying species’ ability to conserve their niches will be a critical step towards determining which species are most likely to adapt to increasing short-term variability in climate.

## Acknowledgements

We thank J. Makinen and M. Lu for their modeling advice and thoughtful comments on the analyses and J. Wilshire, A. Ranipeta, R. Li, and the Map of Life team at the Yale Center for Biodiversity and Global Change for implementing the Spatiotemporal Observation Annotation Tool to annotate occurrence data. We thank F. La Sorte for providing species-level migration distances. We acknowledge funding from NSF grant DEB-1441737 and NASA grants 80NSSC17K0282 and 80NSSC18K0435.

## Data Sharing and Accessibility

GBIF data is available for public use online at https://www.gbif.org/. Code generated to conduct the analyses will be made available in a public repository such as Dryad or Figshare.

## Supporting Information

**Figure S1.**
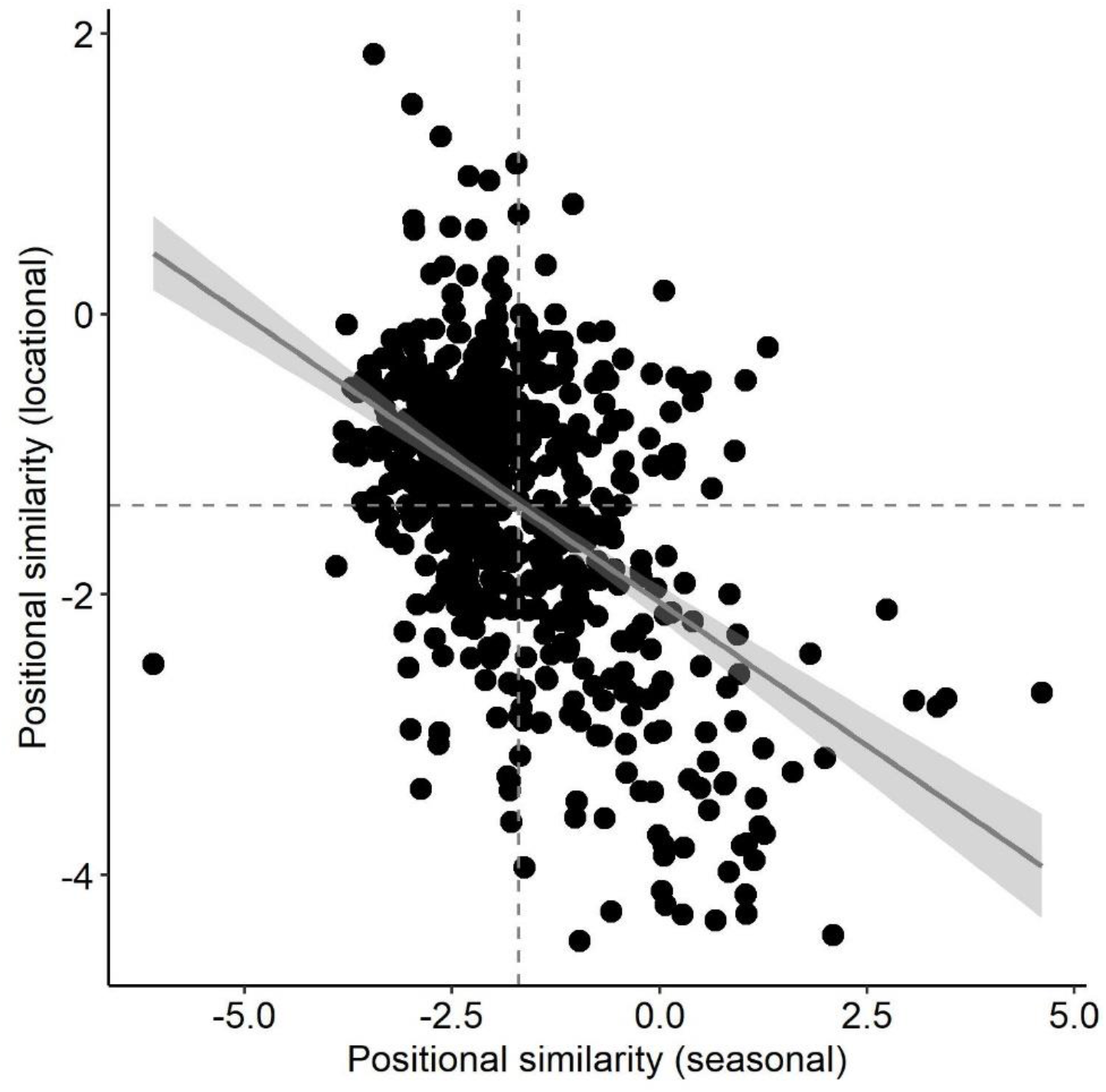
Seasonal and locational niche centroid similarity are inversely related. Niche-tracking species maximize niche similarity between breeding and nonbreeding seasons (defined as seasonal similarity, x-axis) and minimize the similarity between the nonbreeding season niche and the stationary winter niche (defined as locational similarity, y-axis). The reverse is true for non-tracker species. Tropical residents are an exception to this rule; these species have overlapping niche space for all three components, thus maximizing both seasonal and locational similarity. Dashed lines represent means and gray shading represents 95% confidence interval for trend line.

**Figure S2:**
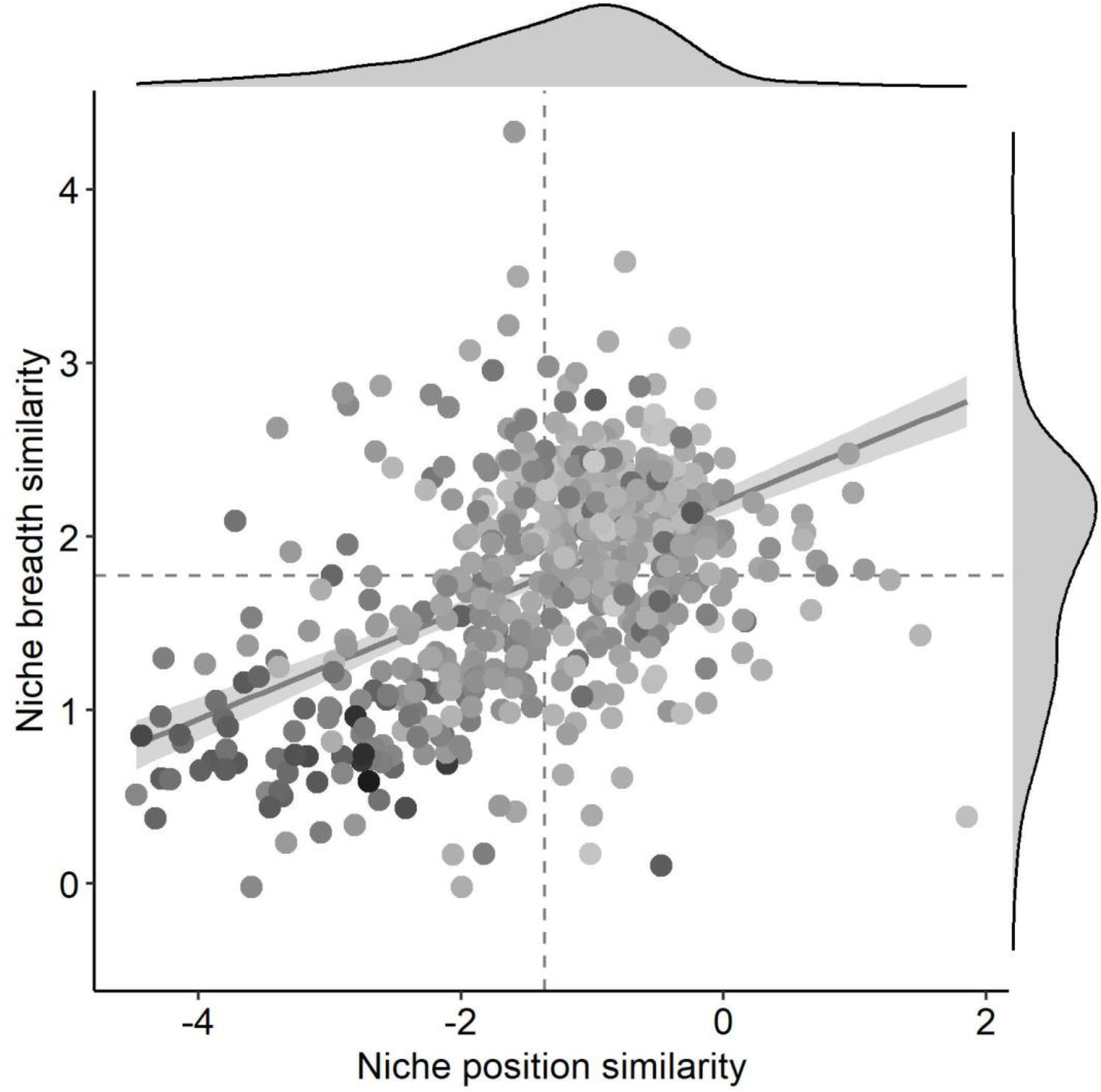
Interspecific variation in locational niche tracking of niche position and breadth. Locational niche similarity represents the similarity of the overwintering niche to the winter conditions at breeding locations. Points represent species-level variation in locational similarity of niche position and breadth between the breeding and overwintering seasons for 619 species of North American birds. Marginal density plots illustrate a positive relationship between tracking locational niche position and breadth. Darker points reflect greater values of seasonal positional niche similarity and are primarily distributed among points with lower values of locational positional similarity.

## Appendix 1

Seasonal niche dissimilarity for 619 species of North American birds (sorted taxonomically). Dissimilarity is partitioned into Mahalanobis distance and determinant ratio for both seasonal (breeding/overwintering comparison) and locational (overwintering/winter at breeding sites comparison) dissimilarity. Position and breadth similarity (visualized and used in analyses) was generated from these values by ln-transforming and sign-flipping.

**(appendix_1.csv)**

## Appendix 2

Updates to avian phylogeny. We updated the avian phylogeny of Jetz et al (2012) to the 2021 Clements taxonomy by harmonizing species names, as follows.

**(appendix_2.csv)**

**Table S1.** Model tables (pgls and ANOVA) summarizing functional trait drivers of seasonal niche position and breadth similarity across species in multivariate models.

| Mahalanobis distance |  |  |  |  |
| --- | --- | --- | --- | --- |
| PGLS table | Coefficient | SE | t-value | p-value |
| Intercept | -1.745 | 0.270 | -6.472 | <0.001 |
| Migration distance | 0.428 | 0.041 | 10.359 | <0.001 |
| Body mass (log) | -0.798 | 0.097 | -8.233 | <0.001 |
| Diet (herbivore) | -0.647 | 0.178 | -3.626 | <0.001 |
| Diet (insectivore) | -0.048 | 0.177 | -0.275 | 0.784 |
| Diet (omnivore) | -0.394 | 0.139 | -2.836 | 0.005 |
| Habitat (generalist) | 0.001 | 0.174 | 0.005 | 0.996 |
| Habitat (open) | -0.381 | 0.110 | -3.470 | 0.001 |
| Habitat (water) | 0.364 | 0.172 | 2.110 | 0.035 |

| Mahalanobis distance |  |  |  |
| --- | --- | --- | --- |
| ANOVA table | DF | F-value | p-value |
| Intercept | 1 | 121.713 | <0.001 |
| Migration distance | 1 | 197.317 | <0.001 |
| Body mass (log) | 1 | 93.094 | <0.001 |
| Diet | 3 | 5.702 | 0.001 |
| Habitat preference | 3 | 9.675 | <0.001 |

| Determinant ratio |  |  |  |  |
| --- | --- | --- | --- | --- |
| PGLS table | Coefficient | SE | t-value | p-value |
| Intercept | 1.360 | 0.262 | 5.183 | <0.001 |
| Migration distance | 0.250 | 0.040 | 6.212 | <0.001 |
| Body mass (log) | 0.525 | 0.094 | 5.565 | <0.001 |
| Diet (herbivore) | -0.053 | 0.174 | -0.305 | 0.760 |
| Diet (insectivore) | 0.206 | 0.172 | 1.196 | 0.232 |
| Diet (omnivore) | 0.102 | 0.135 | 0.754 | 0.451 |
| Habitat (generalist) | -0.081 | 0.169 | -0.481 | 0.631 |
| Habitat (open) | -0.400 | 0.107 | -3.738 | <0.001 |
| Habitat (water) | 0.319 | 0.168 | 1.905 | 0.057 |

| Determinant ratio |  |  |  |
| --- | --- | --- | --- |
| ANOVA table | DF | F-value | p-value |
| Intercept | 1 | 185.261 | <0.001 |
| Migration distance | 1 | 37.702 | <0.001 |
| Body mass (log) | 1 | 51.669 | <0.001 |
| Diet | 3 | 2.201 | 0.087 |
| Habitat preference | 3 | 10.145 | <0.001 |

**Table S2.** Model tables (pgls and ANOVA) summarizing functional trait drivers of locational niche position similarity across species in multivariate models.

| Mahalonobis distance |  |  |  |  |
| --- | --- | --- | --- | --- |
| PGLS table | Coefficient | SE | t-value | p-value |
| Intercept | 0.182 | 0.282 | 0.645 | 0.519 |
| Migration distance | -0.331 | 0.043 | -7.660 | <0.001 |
| Body mass (log) | 0.952 | 0.101 | 9.389 | <0.001 |
| Diet (herbivore) | -0.215 | 0.187 | -1.155 | 0.249 |
| Diet (insectivore) | -0.176 | 0.185 | -0.956 | 0.340 |
| Diet (omnivore) | -0.092 | 0.145 | -0.637 | 0.524 |
| Habitat (generalist) | 0.822 | 0.182 | 4.519 | <0.001 |
| Habitat (open) | 0.415 | 0.115 | 3.609 | <0.001 |
| Habitat (water) | 0.515 | 0.180 | 2.861 | 0.004 |

| Mahalonobis distance |  |  |  |
| --- | --- | --- | --- |
| ANOVA table | DF | F-value | p-value |
| Intercept | 1 | 11.162 | 0.001 |
| Migration distance | 1 | 111.730 | <0.001 |
| Body mass (log) | 1 | 111.154 | <0.001 |
| Diet | 3 | 0.703 | 0.551 |
| Habitat preference | 3 | 7.757 | <0.001 |

**Table S3.** Model tables (pgls) summarizing functional trait drivers of seasonal niche position and breadth similarity across species in univariate models.

| Mahalonobis distance |  |  |  |  |
| --- | --- | --- | --- | --- |
| PGLS table | Coefficient | SE | t-value | p-value |
| Intercept | -3.060 | 0.214 | -14.291 | 0.000 |
| Migration distance | 0.539 | 0.042 | 12.747 | 0.000 |
| PGLS table | Coefficient | SE | t-value | p-value |
| Intercept | -1.039 | 0.219 | -4.744 | 0.000 |
| Body mass (log) | -1.057 | 0.097 | -10.883 | 0.000 |
| PGLS table | Coefficient | SE | t-value | p-value |
| (Intercept) | -1.254 | 0.257 | -4.876 | 0.000 |
| Diet (herbivore) | -1.287 | 0.202 | -6.380 | 0.000 |
| Diet (insectivore) | 0.085 | 0.208 | 0.410 | 0.682 |
| Diet (omnivore) | -0.681 | 0.155 | -4.409 | 0.000 |
| PGLS table | Coefficient | SE | t-value | p-value |
| Intercept | -1.864 | 0.260 | -7.168 | 0.000 |
| Habitat (generalist) | -0.356 | 0.211 | -1.684 | 0.093 |
| Habitat (open) | -0.375 | 0.134 | -2.796 | 0.005 |
| Habitat (water) | 0.121 | 0.208 | 0.578 | 0.563 |

| Determinant ratio PGLS |  |  |  |  |
| --- | --- | --- | --- | --- |
| table | Coefficient | SE | t-value | p-value |
| Intercept | 1.891 | 0.201 | 9.393 | <0.001 |
| Migration distance | 0.229 | 0.040 | 5.770 | <0.001 |
| table | Coefficient | SE | t-value | p-value |
| Intercept | 1.938 | 0.201 | 9.659 | <0.001 |
| Body mass (log) | 0.480 | 0.089 | 5.397 | <0.001 |
| table | Coefficient | SE | t-value | p-value |
| (Intercept) | 2.092 | 0.227 | 9.213 | <0.001 |
| Diet (herbivore) | 0.071 | 0.178 | 0.401 | 0.688 |
| Diet (insectivore) | 0.354 | 0.183 | 1.934 | 0.054 |
| Diet (omnivore) | 0.435 | 0.136 | 3.190 | 0.001 |
| table | Coefficient | SE | t-value | p-value |
| Intercept | 2.304 | 0.218 | 10.572 | <0.001 |
| Habitat (generalist) | -0.161 | 0.177 | -0.909 | 0.364 |
| Habitat (open) | -0.437 | 0.113 | -3.881 | <0.001 |
| Habitat (water) | 0.499 | 0.175 | 2.858 | 0.004 |

**Table S4.** Model tables (pgls) summarizing functional trait drivers of locational niche position similarity across species in univariate models.

| Mahalonobis |  |  |  |  |
| --- | --- | --- | --- | --- |
| distance PGLS table | Coefficient | SE | t-value | p-value |
| Intercept | 1.485 | 0.224 | 6.643 | <0.001 |
| Migration distance | -0.424 | 0.044 | -9.610 | <0.001 |
| distance PGLS table | Coefficient | SE | t-value | p-value |
| Intercept | -0.357 | 0.215 | -1.658 | 0.098 |
| Body mass (log) | 1.119 | 0.095 | 11.721 | <0.001 |
| distance PGLS table | Coefficient | SE | t-value | p-value |
| (Intercept) | 0.331 | 0.265 | 1.252 | 0.211 |
| Diet (herbivore) | 0.521 | 0.208 | 2.510 | 0.012 |
| Diet (insectivore) | -0.255 | 0.214 | -1.194 | 0.233 |
| Diet (omnivore) | 0.351 | 0.159 | 2.207 | 0.028 |
| distance PGLS table | Coefficient | SE | t-value | p-value |
| Intercept | 0.091 | 0.256 | 0.354 | 0.723 |
| Habitat (generalist) | 1.068 | 0.208 | 5.124 | <0.001 |
| Habitat (open) | 0.408 | 0.132 | 3.085 | 0.002 |
| Habitat (water) | 0.774 | 0.205 | 3.770 | <0.001 |

**Table S5.**
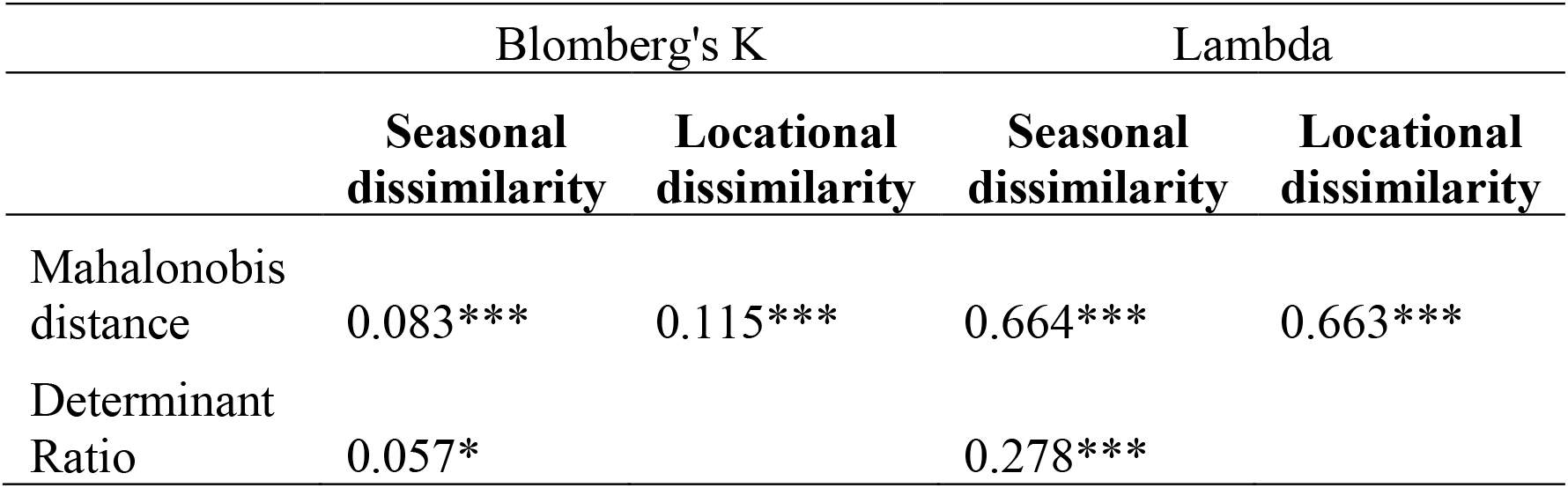
Metrics describing phylogenetic signal in seasonal niche tracking. Asterisks denote significance level (*** < 0.001 < ** < 0.01 < * < 0.05).

